# Biological versus Technical Reliability of Epigenetic Clocks and Implications for Disease Prognosis and Intervention Response

**DOI:** 10.1101/2025.10.13.682176

**Authors:** Raghav Sehgal, Daniel Borrus, John Gonzalez, Yaroslav Markov, ADNI, Albert Higgins-Chen

## Abstract

DNA methylation–based aging biomarkers, or epigenetic clocks, are increasingly used to estimate biological age and predict health outcomes. Their translational utility, however, depends not only on predictive accuracy but also on reliability, the ability to provide consistent results across technical replicates and repeated biological measures. Here, we leveraged the TranslAGE platform to comprehensively evaluate the technical and biological reliability of 18 Epigenetic clocks, including chronological predictors, mortality predictors, pace-of-aging measures, reliable variants, and newer explainable clocks. Technical reliability was quantified across four independent datasets. For standard replicate assays on EPIC and 450K arrays, nearly all clocks achieved excellent technical reproducibility. However, some clocks showed dramatic drops in technical reliability based on differences in slide position and DNA extraction protocol. PC-based clocks, especially PCGrimAge and SystemsAge remained technically reliable in all cases. In contrast, biological reliability, measured across repeated samples collected within hours, before and after meals, under acute stress, across environmental exposures, and over days, was markedly lower, with most clocks showing only moderate stability. PCGrimAge was the only clock with good ICC > 0.75 for biological reliability. Importantly, technical reproducibility did not predict biological reliability; clocks that were technically robust often proved biologically unreliable. We further demonstrated that reliability directly constrains downstream applications. Clocks with higher ICCs produced more stable prognostic associations with cognitive decline and more consistent responsiveness to a vegan diet intervention, whereas unreliable clocks yielded highly variable or spurious effects. Together, these findings reveal that technical reliability is not enough: biological reliability remains a critical limitation for many DNA methylation clocks that constrains their utility, and our work provides a roadmap for prioritizing next-generation clocks most suited for clinical translation.

## Introduction

DNA methylation–based (DNAm) aging biomarkers, widely known as epigenetic clocks, have become leading candidates for quantifying biological age.^1–3^ These measures predict mortality, morbidity, and a range of health outcomes with stronger associations than chronological age alone.^4–6^ Because they can be derived from a simple blood draw, DNAm biomarkers hold substantial promise for research, clinical trials, and eventually routine clinical care.

However, despite their rapid adoption, DNA methylation biomarkers face a critical bottleneck: reliability.^7–10^ For any biomarker to be clinically useful for longitudinal tracking, it must provide consistent and interpretable results across both repeated technical measurements and repeated biological samples.^11,12^ We previously showed that bolstering technical reliability of epigenetic clocks can increase power and reduce false positives in intervention studies.^7,13^

Two forms of reliability are essential to distinguish. Technical reliability reflects reproducibility across test–retest measurements from the same blood sample. Biological reliability reflects stability across repeated collections from the same individual across short time frames (hours or days).^11,12,14–17^ During this interval, events of daily living such as meals, normal levels of stress, circadian variation, or minor environmental exposures may occur. Prior work has demonstrated that first and second generation clocks suffer from large technical variability, sometimes producing deviations of nearly a decade between replicates.^6,7^ Newer clocks such as the PC-based clocks and DunedinPACE were explicitly designed to improve test–retest reproducibility, but remains unclear whether such clocks also necessarily achieve greater biological reliability.

This distinction is not trivial. A clock may be technically precise but biologically unstable, yielding values that fluctuate widely in response to acute, reversible, and minor exposures.^18,19^ Such instability risks undermining their clinical and research utility and lead to unexpected sources of confounding, particularly for longitudinal studies. For example, if a meal can dramatically impact a score, then an intervention study that did not control for fasting status pre- and post-intervention may detect changes that are mistakenly attributed to the intervention but actually reflect trivial changes due to meals. Importantly, the impact of reliability on downstream performance, namely whether clocks with higher ICCs^20^ produce more reproducible prognostic associations and responsiveness estimates, has not been systematically quantified.

Here, we address these gaps by benchmarking the technical and biological reliability of major DNAm clocks spanning five generations: early chronological age predictors (Gen 1), outcome-trained models (Gen 2), pace-of-aging measures (Gen 3), reliable variants of earlier clocks (Gen R), and mechanistically motivated explainable clocks (GenX). These GenX clocks include SystemsAge, OMICmAge, and CausalAge ^21–23^. To perform this analysis in a harmonized manner across clocks and datasets, we utilize our TranslAGE platform which we previously used to systematically report the effects of 51 different interventions^24^. Using replicate data, we quantify technical ICCs across arrays, slide positions, and extraction protocols. To probe biological reliability, we test clock stability under short-term perturbations including meals, stress, pollution exposure, and altitude change. Finally, we directly examine how reliability constrains downstream applications by linking ICCs to the reproducibility of prognostic associations with future cognitive decline and responsiveness to interventions such as a vegan diet. This comprehensive analysis reveals that while most clocks achieve excellent technical reliability, their biological instability limits prognostic and interventional utility, identifying reliability as the central determinant for translating DNAm biomarkers into clinical practice.

## Results

### Variations in Slide Placement and DNA Extraction Reduces Technical Reliability for Some Clocks

We evaluated the technical reproducibility of a broad set of epigenetic clocks across four independent datasets, encompassing replicate samples processed under different laboratory conditions (Figure 1). Overall, the pooled analysis showed that the majority of clocks achieved excellent technical reliability (ICC > 0.9), with SystemsAge, GrimAge, and PC-based derivatives consistently ranking among the most stable. Earlier-generation clocks such as Hannum, PhenoAge, and DNAm PhenoAge displayed relatively lower but still acceptable levels of reliability, generally in the “good” range (ICC ∼0.7–0.8).

**Figure 1.**
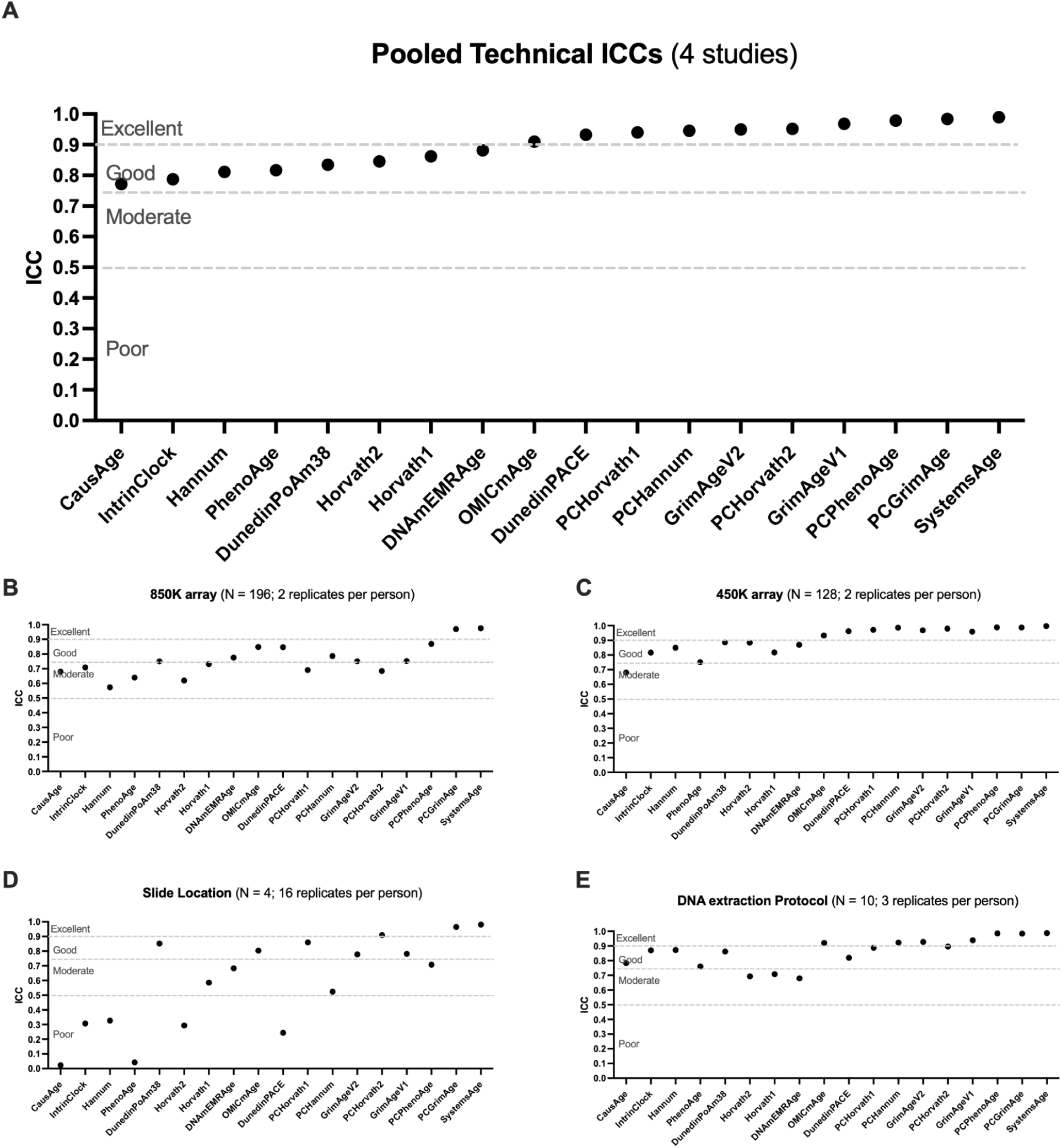
Technical reliability (ICC) of epigenetic clocks across datasets and experimental factors. (A) Pooled technical intraclass correlations (ICCs) for 4 independent studies demonstrate the relative reliability of 18 commonly used epigenetic clocks. Dashed lines denote conventional reliability thresholds (Excellent ≥ 0.90, Good ≥ 0.75, Moderate ≥ 0.50, Poor < 0.50). SystemsAge and principal-component (PC) versions of established clocks show the highest technical reproducibility. (B–E) Stratified ICCs highlight variation across experimental conditions: (B) 850K EPIC array (N = 196; 2 replicates per person), (C) 450K array (N = 128; 2 replicates per person), (D) slide location effects (N = 4; 16 replicates per person), and (E) DNA extraction protocols (N = 10; 3 replicates per person). Across all factors, PC-based measures demonstrate superior reliability, whereas Gen 1 models show greater variability across array type and processing conditions.

When stratified by array platform, both the Illumina 850K (N = 196; 2 replicates per person) and 450K (N = 128; 2 replicates per person) arrays demonstrated high reproducibility across most clocks, reinforcing that array type alone does not substantially compromise technical stability. However, further sensitivity analyses revealed notable drops in reliability under specific laboratory conditions. In particular, clock estimates exhibited marked variability when replicates differed by slide location (N = 4; 16 replicates per person), where several clocks fell below the threshold of “good” reliability and some approached only “moderate” levels. Similarly, while the majority of clocks remained stable under different DNA extraction protocols (N = 10; 3 replicates per person), a subset displayed diminished reproducibility, highlighting that upstream sample processing can introduce additional noise.

Taken together, these results demonstrate that while epigenetic clocks are generally technically robust across arrays and replicates, their performance can degrade under more subtle laboratory perturbations such as slide placement and DNA extraction method. These findings emphasize the need for careful attention to laboratory workflows when applying clocks in clinical or research settings, while also suggesting that biological rather than technical variation likely accounts for much of the observed instability in downstream prognostic and interventional applications.

### Short-Term Exposures Reduce Biological Reliability of Epigenetic Clocks

In contrast to the consistently high technical reproducibility of epigenetic clocks, their *biological reliability* across repeated measures within the same individuals was substantially lower after single short-term events such as meals, stress, pollution, and altitude change (Figure 2). We first tested for systematic changes up or down for the clocks, and found that only altitude change after 7 days caused a systematic shift. We removed time points after this period for altitude change, so that remaining analyses would reveal only random biological noise shifts. Pooled estimates from four independent studies revealed that most clocks achieved only “moderate” to “good” reliability (ICC ≈ 0.4–0.7), with no clock reaching the “excellent” range. Clocks such as GrimAge (versions I and II), DNAmEMRAge, and DunedinPACE generally clustered near the lower end of this spectrum, while PC-based and later-generation clocks (e.g., PCGrimAge, SystemsAge, OMICmAge) performed somewhat better but still fell short of the reproducibility observed in technical replicates.

**Figure 2.**
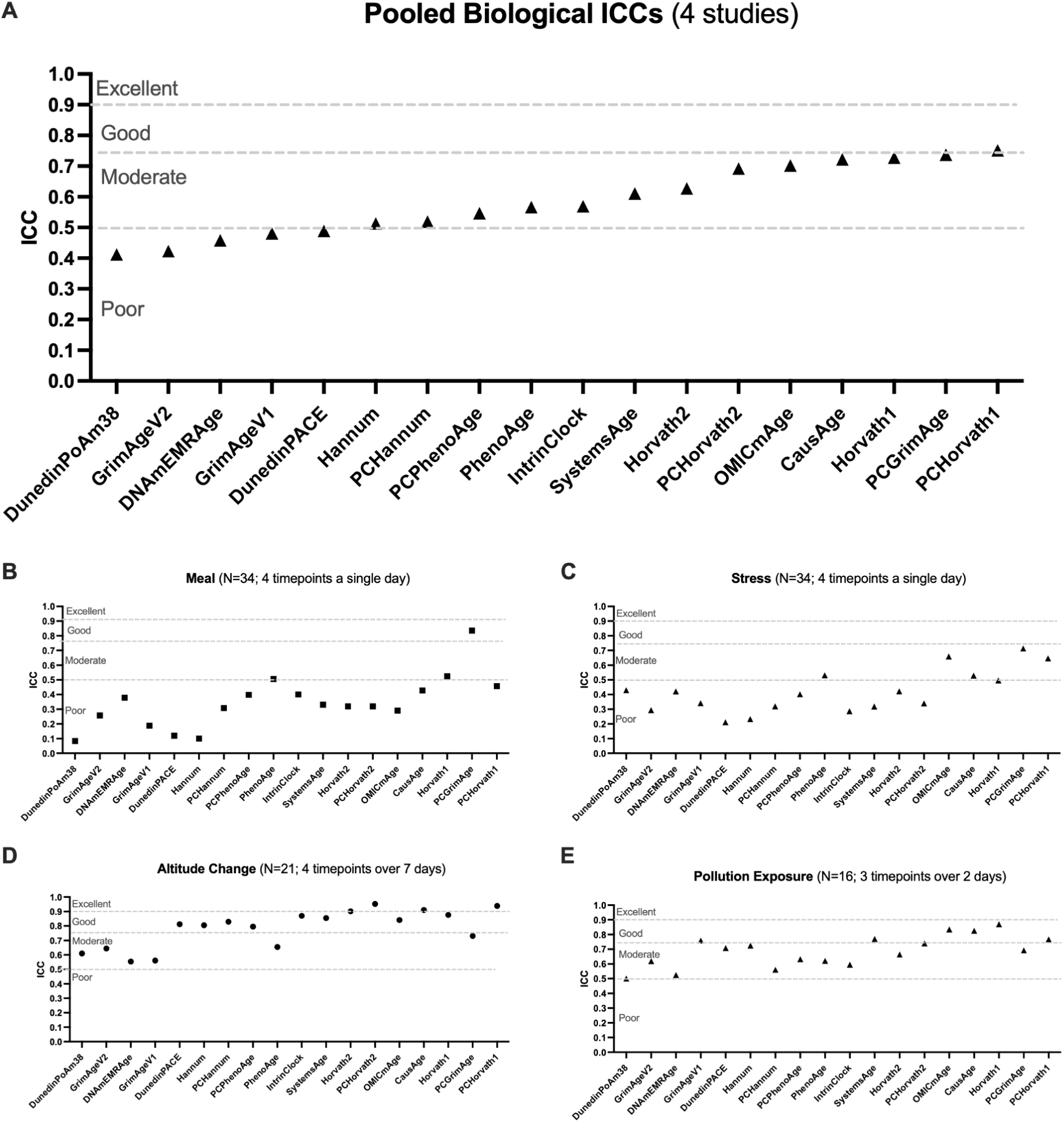
Biological reliability (ICC) of epigenetic clocks across short-term physiological perturbations. (A) Pooled biological intraclass correlations (ICCs) from four independent studies measuring repeated biological samples indicate variable short-term reproducibility among 18 epigenetic clocks. Dashed lines denote standard ICC reliability thresholds (Excellent ≥ 0.90, Good ≥ 0.75, Moderate ≥ 0.50, Poor < 0.50). Multi-system and principal-component (PC) clocks such as SystemsAge, PCPhenoAge, and PCGrimAge show greater biological stability than single-tissue or unadjusted versions. (B–E) Stratified ICCs by acute or environmental challenge illustrate differential within-person biological variability: (B) postprandial effects after a standardized meal (N = 34; 4 timepoints in one day), (C) acute psychosocial stress (N = 34; 4 timepoints in one day), (D) altitude acclimatization (N = 21; 4 timepoints over 7 days), and (E) short-term air pollution exposure (N = 16; 3 timepoints over 2 days). Biological stability was highest under chronic or environmental exposures (altitude, pollution) and lowest under acute perturbations (meal, stress), underscoring the importance of temporal and physiological context in interpreting DNA methylation–based aging biomarkers.

Condition-specific analyses highlighted the sensitivity of clocks to short-term biological fluctuations. Following a single meal (N = 34; 4 timepoints within a day), reliability dropped markedly, with many clocks performing only in the “moderate” or “poor” range, suggesting strong acute biological variability. Stress exposure (N = 34; 4 timepoints within a day) produced a similar effect, with several widely used clocks including DunedinPACE and GrimAge falling into the “poor” reliability range. Pollution exposure (N = 16; 3 timepoints over 2 days) also induced sharp drops in stability, reinforcing the susceptibility of clock estimates to short-term environmental perturbations. By contrast, changes across altitude (N = 21; 7 timepoints over 21 days) showed comparatively higher stability, with several clocks approaching “good” reliability, suggesting that slower or more systemic physiological adaptations may exert less volatility on methylation-based age measures than acute perturbations.

Together, these findings demonstrate that while epigenetic clocks are technically robust, their biological instability under everyday exposures: including diet, stress, and environmental conditions, poses a significant limitation.

### Technical Reliability does not track with Biological Reliability of Epigenetic Clocks

We examined the relationship between technical and biological reliability across clocks (Figure 3 A). While nearly all clocks achieved high technical ICCs, their corresponding biological ICCs were substantially lower, with no clear correlation between the two metrics (r = 0.0168). For example, SystemsAge and PC-based clocks (e.g., PCGrimAge, PCHorvath1/2) were among the most technically stable and showed comparatively higher biological reliability, yet widely used measures such as GrimAgeV2 and DunedinPACE remained technically excellent but biologically unreliable.

**Figure 3.**
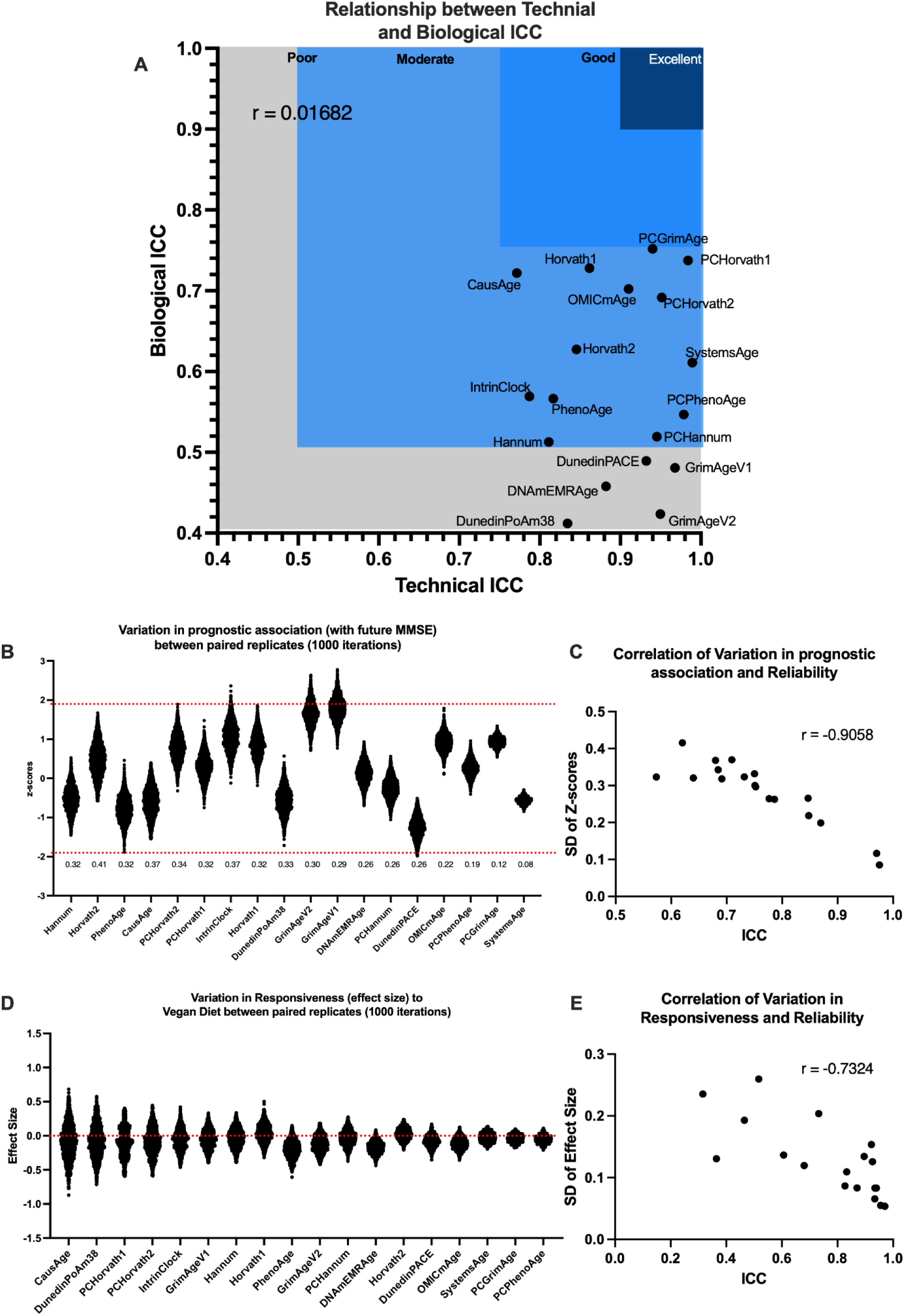
Relationship between technical reliability, biological stability, and downstream analytic variability of epigenetic clocks. (A) Relationship between pooled technical and biological intraclass correlations (ICCs) across 17 epigenetic clocks demonstrates that technical and biological reproducibility are largely independent (r = 0.0168). Principal-component (PC) clocks (e.g., PCPhenoAge, PCGrimAge, PCHorvath1, Systems Age) show high reproducibility in both dimensions, while first-generation clocks and Dunedin-based measures exhibit lower stability. (B) Variation in prognostic associations with future Mini-Mental State Examination (MMSE) scores across 1,000 resampling iterations illustrates greater variability for lower-ICC clocks. (C) The standard deviation of prognostic Z-scores is strongly inversely correlated with reliability (r = –0.9058), indicating that clocks with higher ICCs yield more consistent effect estimates across replicates. (D) Variation in responsiveness (effect size) to a vegan diet intervention across 1,000 paired resampling iterations reveals that unreliable clocks show greater dispersion in estimated effects. (E) Standard deviation of responsiveness effect sizes is also inversely correlated with ICC (r = –0.7324). Together, these analyses show that clocks with higher technical and biological reliability produce more stable and reproducible estimates of both prognostic and interventional outcomes.

### Clock instability undermines Prognostic Power and Intervention Responsiveness

We next evaluated how clock reliability shapes their utility in prognostic and interventional settings (Figure 3 B-D). Using repeated resampling of paired technical replicates (1,000 iterations), we quantified the stability of associations between clocks and two key outcomes: (1) prognostic prediction of future cognition, measured by Mini-Mental State Examination (MMSE), ^25^ and (2) responsiveness to intervention, estimated as effect sizes for a vegan diet exposure.

For prognosis, clocks with lower reliability displayed strikingly wide variability in their associations with future MMSE scores. Across replicates, low-ICC clocks such as Hannum, PhenoAge, and DNAmEMRAge produced highly unstable effect estimates, with z-scores fluctuating across the full significance spectrum. In contrast, high-reliability clocks including SystemsAge, PC-based GrimAge, and PCPhenoAge produced far narrower distributions centered around consistent effect sizes. Quantitatively, reliability strongly predicted the stability of prognostic performance: the standard deviation of z-scores across replicates correlated negatively with Technical ICC (r = –0.91), demonstrating that clocks with higher reliability yield reproducible prognostic signals, while biologically unstable clocks may generate spurious or inconsistent associations.

A parallel pattern was observed for responsiveness. When estimating the impact of vegan diet exposure, less reliable clocks showed large variability in effect sizes across replicates, often flipping direction or yielding inflated associations. In contrast, technically and biologically stable clocks maintained consistent effect size estimates, even under repeated resampling. The correlation between Technical ICC and variability in responsiveness (r = –0.73) confirmed that reliability is a strong determinant of whether a clock can robustly capture intervention effects.

Together, these analyses establish reliability as a critical prerequisite for the translational application of epigenetic clocks. Clocks that are technically robust but biologically unstable can undermine both prognostic predictions and responsiveness assessments, leading to misleading conclusions in clinical or research contexts. Conversely, clocks with higher reliability offer stable, reproducible signals that are more suitable for use in longitudinal studies and interventional trials.

## Discussion

The promise of epigenetic clocks lies in their ability to act as prognostic and interventional biomarkers, yet our results show that their utility is critically constrained by reliability.^7,26^ While discussions of biomarkers often emphasize accuracy and predictive value, the consistency of measurement across both technical and biological contexts is equally essential for clinical translation and longitudinal studies.^26^ This study provides the most comprehensive evaluation to date of both technical and biological reliability across multiple generations of DNA methylation (DNAm) clocks, and directly demonstrates how reliability governs prognostic and interventional performance.

Our findings confirm that technical reliability is largely a solved problem. Across four independent datasets, most clocks demonstrated excellent reproducibility in replicate assays, with PC-based clocks such as SystemsAge consistently ranking among the most stable. Technical robustness also translated into greater consistency in prognostic predictions, as shown by narrower distributions of associations with cognitive decline and brain volume loss. These results underscore that technical noise, while once a significant limitation for early clocks such as Horvath1, Hannum, and PhenoAge, has been effectively addressed in Generation Reliable models.

By contrast, biological reliability remains a major obstacle. We found that repeated measures taken under common short-term perturbations, including meals, stress, and pollution exposure,caused substantial fluctuations in epigenetic age estimates, with most clocks falling into only moderate reliability at best.^7,27,28^ Critically, technical reproducibility did not predict biological stability: some of the most technically robust clocks, such as GrimAgeV2 and DunedinPACE, were among the most biologically fragile. This dissociation reveals that laboratory precision alone is not sufficient to ensure a biomarker’s clinical utility.

The consequences of biological instability are not merely theoretical. Our analyses show that clocks with lower reliability produce unstable prognostic predictions and inconsistent intervention responsiveness. Associations with cognitive decline and dietary exposures varied widely across replicate analyses for unreliable clocks, raising the risk of spurious or misleading findings. We previously warned about false positive intervention results in clocks with low reliability, especially chronological age clocks ^13,24^. Conversely, clocks with higher reliability, particularly PCGrimAge and SystemsAge, provided stable signals across both prognostic and interventional contexts. These results highlight reliability as the central determinant of translational readiness. Importantly, there was no relationship between technical reproducibility and biological stability of clocks, reinforcing the importance of distinguishing between laboratory precision and biological robustness when evaluating clock performance.

It is worth mentioning that these results should be taken in context of other available biomarkers. Even though we see moderate biological reliability from the best Generation Reliable DNAm biomarkers such as PCGrimAge (ICC = 0.76), they are still better or comparable to several well-known biomarkers currently in clinical practice or used as surrogate endpoints. For example, Aβ40 shows a time-of-day varying ICC of 0.76, Aβ42 an ICC of 0.88, and GFAP an ICC of 0.83.^29^ Clinical biomarkers like hematocrit have been reported with ICCs as low as 0.61 pre- and post-hydration. ^30^ Moreover, meal consumption or fasting can alter protein biomarkers such as insulin (ICC = 0.75), C-peptide (ICC = 0.66), and free IGF-1 (ICC = 0.55) ^31^. Other widely used proteomic biomarkers show even lower stability in response to perturbations such as exercise training, including BDNF (ICC = 0.26), IL-1β (ICC = 0.65), IL-6 (ICC = 0.0), and IFN-γ (ICC = 0.11)^32^. These comparisons suggest that while DNAm biomarkers are biologically unstable, their reliability is not unusually poor when judged against other biomarker types, and in some cases DNAm biomarkers are superior.

This study has several limitations. First, while we examined reliability across multiple datasets and perturbations, our analyses remain constrained to specific populations and conditions. Larger, more diverse cohorts are needed to test whether the observed patterns generalize across ancestries, age groups, and disease states. Second, some may argue that the biomarker changes we observe after events of daily life such as meals, stress, and altitude change may be bona fide changes in the aging process. However, we do not think it is plausible that such minor single events should truly age or de-age a person by multiple years. Even if these events did have such a large effect, it still highlights the importance of identifying methods to separate their effects from intervention treatment effects occurring in the same period. Third, the biological perturbations we tested were relatively short-term and acute. Longitudinal studies spanning years are required to determine whether biological instability resolves over longer intervals or reflects intrinsic noise in methylation dynamics. Fourth, while we identified correlations between reliability and prognostic or interventional stability, causal mechanisms underlying biological instability remain unclear. Immune cell dynamics, stress-related pathways, and metabolic fluctuations are likely contributors, but further mechanistic studies will be necessary.^33^ Finally, although we focused on widely used clocks, the field is rapidly evolving, and newer models may address some of the limitations highlighted here.

In conclusion, our findings demonstrate that while epigenetic clocks now achieve excellent technical reproducibility, biological instability remains the central limitation to their clinical translation. Reliability is the key constraint shaping prognostic and interventional utility, and efforts to develop next-generation biomarkers must prioritize not only technical robustness but also resilience to biological variability.

## Data Availability Statement

All ICC values and related values will be available upon publication. Code to calculate all clocks except for OMICmAge and DNAmEMRAge are accessible at https://github.com/HigginsChenLab/methylCIPHER. Code to calculate OMICmAge, DNAmEMRAge, and underlying algorithms will be accessible via TruDiagnostic’s DNAm Analysis Software after publication. You can request access to the software at https://www.trudiagnostic.com/softwarerequest. We are currently working on a platform to download all TranslAGE harmonized data drawn from public sources.

## Acknowledgments

This work was supported by the National Institute on Aging (NIA:1R01AG065403 to A.H.C.). It was also supported by the Impetus Grant (R.S.), the Gruber Science Fellowship at Yale University (R.S.), and the Thomas P. Detre Fellowship Award in Translational Neuroscience Research from Yale University (to A.H.C.).

## Author Contributions

R.S., D.S.B. and A.H.C. conceived the project. R.S. conceived the study design. R.Sehgal performed reliability analysis. R.S., D.S.B., and A.H.C. built the pipeline for TranslAGE. R.S. performed study data and metadata curation for reliability datasets. Y.M. processed raw ADNI DNA methylation data. R.S, D.S.B., and A.H.C. wrote the manuscript, and all authors reviewed and contributed to the manuscript.

## Methods

### Analysis Pipeline - TranslAGE Reliability

#### Step 1: Data Curation

We began by compiling a comprehensive list of publicly available reliability datasets. Publicly available datasets were sourced from Gene Expression Omnibus (GEO) and the European Molecular Biology Laboratory (EMBL) repositories and other public data repositories. Datasets were selected on the basis of the following criterion: 1) DNAm data is either for technical or biological replicates, 2) Tissue of origin for the DNAm data is blood (ideally PBMCs), 3) Study passes quality checks. For each study, we curated extensive study-level metadata, including the following key variables:

● **Type of technical or biological variability**: Identifying the different possible variations that could be possible given a set of replicate samples.
● **Demographic data**: Capturing demographic distribution to test reliability across a wide gamut of age, sex and other demographic parameters.

These metadata were manually extracted from study documentation and supplementary materials to ensure accuracy. After Careful curation the following datasets were curated for analysis.

| Study ID | Array Type | Type | Individual (Replicates) |
| --- | --- | --- | --- |
| ADNI<sup>35</sup> | EPICv1 | Technical | 196(2) |
| GSE174422<sup>36</sup> | 450k | Technical | 128 (2) |
| GSE227809 - Stress <sup>18</sup> | EPICv1 | Biological | 34 (4) |
| GSE227809<sup>18</sup> | EPICv1 | Biological | 34 (4) |
| GSE105123<sup>37</sup> | 450k | Biological | 21(4) |
| GSE56553<sup>38</sup> | 450k | Biological | 16(3) |
| GSE250556<sup>39</sup> | EPICv1 | Technical | 4(16) |
| Zenodo1285774<sup>40</sup> | 450k | Technical | 10(3) |

#### Step 2: Metadata standardization

We utilized a standardized cleaning script that was adapted for each dataset. We standardized the variables across all studies to create uniform, comparable datasets. This harmonization process involved:

● **Standardized columns**: We ensured that all datasets contained consistent variables, including **Age**, **Sex**, **Sample ID**, **Individual ID**, **Replicate condition**. This step allowed for consistent cross-study analysis.
● **Sample identification**: To maintain data integrity, each sample was assigned a unique **Sample ID** and linked to an **Individual ID**, ensuring that repeated measurements from the same individual could be tracked longitudinally.

This harmonization facilitated subsequent analyses, allowing for robust comparisons between different studies and intervention types.

#### Step 3: Clock Calculation

For each of the curated datasets, we calculated over **18 DNA methylation (DNAm) biomarkers** using the **MethylCIPHER 2.0** package. This package allows for the computation of various methylation-based aging and health biomarkers. These biomarkers were calculated for each sample based on the methylation data available in each study.

#### Step 4: Age-residualized scores

We processed the biomarkers by age-residualizing them in each dataset. Age-residualization was performed to control for the natural variation in DNAm biomarkers due to chronological age. The procedure involved linear regression. We used age as the independent variable and each DNAm biomarker as the dependent variable to calculate the residuals. These age-residuals represent the portion of the biomarker that is not explained by chronological age.

#### Step 5: ICC calculation

We performed Intraclass Correlation Coefficient (ICC) analysis to evaluate the reliability of each epigenetic clock. ICC measures the consistency or reproducibility of measurements, with higher values indicating greater reliability. The analysis was conducted using the irr (0.84.1) package in R, specifically using the `icc()` function to compute the ICC for each clock across the paired samples. Function irr::icc(data, model = “twoway”, type = “agreement”, unit = “single”). We used a p-value cut off of 0.05 to determine non-significant ICCs ^41^.

### ICC Pooling Methodology

ICCs were pooled across multiple datasets using a comprehensive meta-analytical approach with both fixed-effects and random-effects models. The pooling procedure involved the following steps:

● **Fisher’s Z-Transformation**: Individual ICC values were transformed using Fisher’s z-transformation [z = 0.5 × ln((1+ICC)/(1-ICC))] to normalize their distribution and stabilize variance, with values clipped to [-0.999, 0.999] to ensure valid transformations.
● **Weighting Scheme**: Study-specific weights were calculated as the inverse of the variance (1/SE²), where standard error was estimated using SE = 1/√(n-3) based on the effective sample size (minimum of subjects and observations).
● **Random-Effects Model**: The DerSimonian-Laird method was employed to account for between-study heterogeneity, incorporating tau-squared (τ²) estimates of between-study variance into the weighting scheme.

Separate meta-analyses were conducted for biological and technical replicability datasets, with 95% confidence intervals calculated for both fixed and random-effects estimates. Results showed substantial heterogeneity (I² > 50%) for most epigenetic clocks, justifying the use of random-effects models as the primary pooling approach.

### Multiverse analysis to assess effects of technical noise on prognostic associations

To evaluate the impact of technical noise on morbidity associations, we performed a multiverse analysis by simulating various scenarios where one technical replicate from each pair was randomly selected for the association analysis. We performed 1000 such association analyses for each epigenetic clock by randomly selecting one of the two technical replicates for each individual in the ADNI dataset.^42^ The goal was to evaluate how technical noise might affect the variability in prognostic association z-scores across multiple analyses.

1. **Random Selection of Replicates**: For each of the 1000 simulated association analyses, we randomly selected one of the two technical replicates for each individual. This process was repeated independently for each simulation to introduce variability in the input data.
2. **Association Analysis for Each Simulation**: We then performed the association analysis using the selected replicates in each simulation, calculating the z-scores for disease associations with each epigenetic clock.
3. **Comparing Z-Score Distributions**: After completing all 1000 simulations, we compared the distribution of the resulting z-scores for each epigenetic clock. This allowed us to quantify the potential variation in association results due to technical noise.

This approach allowed us to systematically explore how technical variation between replicates could influence disease association outcomes for each clock, providing insight into the robustness of the associations in the presence of technical noise.

To further investigate the variability in disease associations due to technical replicates, we conducted an additional association analysis using the unselected replicates. For each DNAm biomarker, the analysis was performed as follows:

1. **Association Analysis on Unselected Replicates**: We performed an association analysis on the replicate that had not been used in the primary analysis. This provided a set of z-scores for disease associations based on the unselected replicates.
2. **Calculating the Difference in Z-Scores**: We then subtracted the z-score obtained from the mirrored group (i.e., the z-score from the primary replicate analysis) from the z-score of the unselected replicate. The difference between these z-scores provided a measure of how much the association changed depending on the randomly selected set of technical replicates.
3. **Comparison of Z-Score Differences Across Biomarkers**: We compared the distribution of these z-score differences across various DNAm biomarkers to assess the impact of replicate selection on disease association variability. This analysis allowed us to further quantify the influence of technical variation on the association results.

